# *Vibrio aquamarinus* sp. nov., a luminous marine bacteria isolated from the Black Sea

**DOI:** 10.64898/2026.01.16.699947

**Authors:** M.A. Sazykina, S.A. Khrulnova, I.S. Sazykin, E.V. Malysheva, S.M. Rastorguev, A.A. Novikov, A.A. Korzhenkov, M.N. Konopleva, A.A. Kudryavtseva, R.V. Berezov, K.V. Mekhantseva, S.V. Bazhenov, V.B. Shirokov, V.A. Chistyakov, I.V. Manukhov

## Abstract

Three novel bioluminescent bacterial strains, VNB-15^T^, VNB-16 and SChm4, were isolated from water of the Black Sea (Russia) and intestines of the Black Sea horse mackerel. Cells of the isolated strains are motile Gram negative slightly curved rods with single polar flagellum. The temperature range for growth was 10-35°C, the optimum being 20-25°C. The pH range for growth was 6.0-9.0, the optimum being 7.0-8.0. The bacteria were able to grow in the presence of 0.5 to 5.0% NaCl (w/v), the optimum being 1.0-4.0% (w/v). Phylogenetic analysis based on comparison of 16S rRNA sequences shows these strains to have kinship with the species *Vibrio jasicida, Vibrio hyugaensis, Vibrio alginolyticus, Vibrio campbelli, Vibrio rotiferianus, Vibrio harveyi* and *Vibrio owensii* with sequence similarity from 99.6 to 98.0%. Phylogenetic analysis based on comparison of the sequences of genes *gyrB*, *recA*, *pyrH*, *gapA, rpoA, mreB, ftsZ, topA* shows that the strains VNB-15^T^, VNB-16 and SChm4 to form a cluster within the *V. harveyi* clade and belong to a new species of the *Vibrio* genus. Comparison of the complete genomic sequence of VNB-15^T^ with typical strains of nearby species also indicates that VNB-15^T^ belongs to a separate species (maximum similarity 98% with *V. hyugaensis* and 96% with *V. jasicida*). VNB-15^T^ differs from closely related species by its ability to utilize glucose, mannitol, inositol, sorbitol, rhamnose and sucrose, and to form lysine decarboxylase, ornithine decarboxylase, lipase, acid phosphatase, α-glucosidase, β-glucosidase and N-acetyl-β-D-glucosaminidase enzymes. Based on phylogenetic analysis and phenotypic characteristics, *Vibrio aquamarinus* sp. nov. is proposed. The type strain is VNB-15^T^ (= VKPM B-11245^T^ = DSM 26054 ^T^).

**Repositories:** The GenBank accession numbers for the *gapA*, 16S rRNA, *gyrB*, *pyrH, rpoA, recA, mreB, ftsZ, topA* genes sequences of strain VNB-15^T^ are JQ319116-JQ319121, KX242381, KX 242384, KX242387, respectively. The GenBank accession numbers for the *gapA, gyrB*, *pyrH, recA, rpoA*,16S rRNA, *mreB, ftsZ, topA* genes sequences of strain VNB-16 are KP221561-KP221566, KX242382, KX 242385, KX242388 respectively. The GenBank accession numbers for the 16S rRNA, *gapA gyrB, rpoA*, *recA, pyrH, mreB, ftsZ, topA* genes sequences of strain SChm4 are KX242375-KX242380, KX242383, KX 242386, KX242389, respectively. The GenBank/EMBL/DDBJ accession numbers for the housekeeping gene sequences used in this study are detailed in supplementary Table S1, Figures S1-S8. The genome of *Vibrio aquamarinus* sp. nov. VNB-15^T^, comprising two chromosomes and a plasmid, has been assembled and deposited in the NCBI database under the submission number SUB14585067

## Introduction

The *Vibrio* genus comprises dozens of phylogenetically, phenotypically and ecologically isolated species widespread in the environment. Vibrions are among the most common organisms in surface waters all over the world (Farmer et al., 2005; Thompson et al., 2004). Specimens of the *Vibrio* genus are found in marine environments with varying degrees of salinity, in marine sediments, in estuaries. This genus includes strains inhabiting various organisms ranging from plankton to fish. *Vibrio* species live on skin of marine animals and inside the contents of their intestines where they act as facultative symbionts, parasites or as pathogens. Several *Vibrio* species are important causative agents of diseases in marine animals and humans (Maugeri et al., 2000) (Ripabelli et al., 1999).

Constitution of the *Vibrionaceae* family is being regularly clarified and reclassified, with new genera added (Boyd et al., 2015; Urbanczyk et al., 2007; Yoshizawa et al., 2009). Currently, the family *Vibrionaceae* includes the following genera: *Vibrio* (Farmer et al., 2005), *Photobacterium* (Labella et al., 2017), *Salinivibrio* (Mellado et al., 1996), *Grimonita* (Thompson et al. 2003), *Enterovibrio* (Thompson et al., 2002) *Aliivibrio* (Urbanczyk et al., 2007), *Thaumasiovibrio* (Amin et al., 2017) and Candidatus *Photodesmus* (Hendry & Dunlap, 2011).

The majority of the *Vibrio* species are distributed among thirteen phylogenetically robust clades, namely *Anguillarum, Cholerae, Coralliilyticus, Diazotrophicus, Gazogenes, Halioticoli, Harveyi, Nereis, Nigripulchritudo, Orientalis, Scopthalmi, Splendidus* and *Vulnificus*. This distribution is based on the multilocus sequence analysis (MLSA) of nine genes (i.e., *ftsZ*, *gapA*, *gyrB*, *mreB*, *pyrH*, *recA*, *rpoA*, *topA*, and 16S rRNA) (Sawabe et al., 2007). In relatively recent report (Sawabe et al., 2013), eight new clades within the family *Vibrionaceae* (*Damselae*, *Mediterranei*, *Pectenicida*, *Phosphoreum*, *Profundum*, *Porteresiae*, *Rosenbergii*, and *Rumoiensis*) were proposed, in addition to those previously proposed in the report of Sawabe et al. (2007). Molecular techniques allow detection of novel *Vibrio* species, leading to constant updating of its taxonomy (Beaz-Hidalgo et al., 2010, 2010; Cano-Gómez et al., 2010, 2010; Jin et al., 2013, 2013; Lasa et al., 2013; Lucena et al., 2012, 2012; Rameshkumar et al., 2011, 2011).

Several species of *Vibrio* are luminescent. Bioluminescent bacteria are common in the ocean, especially in temperate to warmer waters (Dunlap & Kita-Tsukamoto, 2006). Bioluminescent bacteria of the Black Sea are not studied well. In the literature, their descriptions are scarce, with only a few published works (Katsev, 2002) (Tsybulskii & Sazykina, 2010) (Gretskiĭ, 2014). Low salinity (about 18 permille), chemocline zoning (a layer of dramatic changes in hydrochemical parameters, primarily, the transition between oxygen and hydrogen sulfide zones) and other hydrobiological peculiarities of the Black Sea, along with its relative isolation from the Mediterranean Sea resulted in formation of some habitats specific to this locality and expected to harbor endemic bacterial species. In this paper we attempt to characterize a novel species of luminescent bacteria inhabiting Black Sea waters.

The aim of this study was to determine the taxonomic position of the luminescent strains VNB-15^T^, VNB-16 and SChm4 by means of polyphasic taxonomic approach, including the physiological properties, chemotaxonomic characteristics and phylogenetic analysis based on comparison of sequences of eight housekeeping genes and gene sequences of 16S rRNA.

## Materials and methods

### Isolation and culture conditions

Strains VNB-15^T^ and VNB-16 were isolated from water sampled from the Black Sea in the region of Abrau Durso (44°40′36″ N, 37°33′49″ E) in July 2009 and in June 2014, respectively. Water samples were taken from the surface layer at 100 m distance from the shore. Membrane filter method was utilized to isolate bioluminescent bacteria. Cells were concentrated on membrane filters with pore size of 0.22 microns (“Sartorius AG”, Germany). The filters were then placed on the surface of solid selective media described earlier (Sazykina et al., 2007, 2009). The composition of media was selected while taking into account the salinity of the water in the area of collection (17 %_0_); cells were incubated at 20°C for 18-24 h.

SChm4 strain was isolated from intestines of *Trachurus mediterraneus subsp. ponticus* (Aleev YuG, 1956) caught near the Utes settlement on the coast of southern Crimea (44°59’97²N, 34°37’31²E) in June 2014, then cultivated on SWT medium (5 g/L tryptone, 2.5 g/L yeast extract, 15 g/L sea salt, 10g/L glycerol).

Luminescent colonies grown on the medium in Petri dishes were transferred to fresh sterile medium to isolate a pure culture of bacteria.

All strains were cultured on plates of Marine agar (Difco) at 24±1°C under aerobic conditions. Stock cultures were maintained frozen at −80°C in Marine broth (Difco) supplemented with 15 % glycerol (v/v).

### Phenotypic characterization

Morphological properties, growth and bioluminescence characteristics of bacteria were evaluated at different temperatures; we have also studied their enzymatic properties and ability to ferment various sugars.

The morphology of cells and their flagella was studied using an atomic force microscope (Integra Prima, by NT-MDT). Temperature and pH range for growth of isolated strains were determined by incubating isolated colonies on the Marine agar (Difco).

Phenotypic identification of the three strains VNB-15^T^, VNB-16 and SChm4 and the reference strains followed the methods described by (Alsina & Blanch, 1994; Holt et al., 1994; Lemos et al., 1985; West et al., 1983), that included Gram stain, growth on thiosulphate citrate bile salt sucrose agar (TCBS, Bioxon, Mexico), cell morphology, luminescence, sensitivity to the vibriostatic agent O/129, motility, O–F test, arginine dehydrolase, ornithine decarboxylase and lysine decarboxylase. Salt tolerance (NaCl %: 0, 0.5, 1, 2, 3, 4, 5, 6, 7, 8, 9 and 10), growth at different temperatures (4, 10, 20, 25, 30, 35, 37 and 44°C), growth at different pH (4, 5, 6, 7, 8, 9, 10), indole production, amylase and gelatinase activities, lipase hydrolysis, Voges–Proskauer, and utilization of citrate, L-arabinose were also investigated.

Catalase activity was determined by formation of bubbles in 3 % (v/v) solution of hydrogen peroxide. Oxidase activity was determined by Oxi-test paper strips (Erba Lachema s.r.o.).

Additional phenotypic characteristics were determined using API 20E system (BioMerieux, France), PBDE (biochemical plate) systems (research and manufacturing association “Diagnostic Systems”, Russia), MMTE 1 system of multimicrotests (research and manufacturing association “Allergen”, Russia).

PBDE and MMTE 1 are biochemical plates differentiating enterobacteria. The MMTE 1 system allows performing 12 tests (formation of indole, hydrogen sulfide; presence of lysine decarboxylase, ornithine-decarboxylase, urease, phenylalanine deaminase activities as well as fermentation of mannitol, citrate, malonate, sucrose, lactose and sorbitol). PBDE allows performing 19 tests, including the detection of urease, β-D-galactosidase, lysine decarboxylase, ornithine-decarboxylase and arginine dihydrolase activities; formation of hydrogen sulfide, indole and acetoin; fermentation of glucose, sucrose, mannitol, malonate, citrate, sodium citrate with glucose, inositol, sorbitol, arabinose, maltose).

The media for experiments on the study of phenotypic features contained 1.7 % NaCl. The identification of biochemical parameters was performed by automatic bacteriological analyzer Vitek® 2 Compact 30 (BioMerieux) with cards Vitek® 2 GN (BioMerieux).

### Molecular characteristics

The isolation of the genomic DNA from the cells was carried out as described earlier (Zavilgelsky GB et al., 2002). Sequencing of 16S rRNA gene and eight housekeeping genes was performed by Sanger method. Multilocus sequence analysis (MLSA) of eight concatenated housekeeping gene sequences encoding topoisomerase I (*topA*), cell division protein (*ftsZ*), glyceraldehydes-3-phosphate dehydrogenase (*gapA*), DNA gyrase B subunit (*gyrB*), actin-like cytoskeleton protein (*mreB*), uridylate kinase (*pyrH*), recombination repair protein (*recA*) and RNA polymerase alpha-subunit (*rpoA*) was carried out as described previously (Ast et al., 2009; Sawabe et al., 2007). Manual examination of the sequence chromatograms and their comparisons were carried out in Vector NTI 9 (Informax, USA). Comparisons of individual sequences and their similarity analysis were done using BLAST (http://blast.ncbi.nlm.nih.gov/) and EzTaxon tool (http://eztaxon-e.ezbiocloud.net), Clustal W (https://github.com/yayakri/clustal-project). Phylogenetic relationships among submitted species of the *Vibrio* genus were examined by means of MEGA (Tamura et al., 2021). Phylogenetic trees were constructed by the Neighbor-Joining method. A bootstrap analysis to investigate the stability of the trees was performed in 1000 replicates (accession numbers are listed in Table S1).

The VNB-15T genome was sequenced using nanopore MinION (ONT, UK) with Ligase sequencing kit and Illumina. The quality of the reads was evaluated and visualized using the FastQC program (https://github.com/icaoberg/singularity-fastqc ), after which they were cleaned using Trimomatic (https://github.com/usadellab/Trimmomatic ). Next, the prepared reads were assembled into contigues using the hybrid assembly method in the Flye program (https://github.com/FrancescoSaverioZuppichini/Flue). Next, the sequences were assembled into scaffolds using the ABACAS utility and annotated using Prokka. As a result, two large scaffolds were obtained, and therefore we can assume that these nucleotide sequences are chromosomes and a short contig, which is a plasmid. The rRNA genes were identified using the Barrnap utility (https://github.com/tseemann/barrnap). After that, the assembly was further evaluated using QUAST, which showed the N50 value was 3834711. Verification of the assembly using the CheckM program showed 99.74% completeness and 0.69% contamination, which indicates a high completeness and purity of the genome. Multiple alignment of complete genomes (excluding plasmids) obtained from NCBI was performed using ProgressiveMauve. Based on these alignments, a phylogenetic tree structure was built using FastTree, with branch support calculated from 100 pseudo-loading replicas. The tree was visualized using iTOL.

RelTime-ML method (Tamura et al. 2021) was applied to obtain phylogenetic tree with the dataset of concatenated genes (*gapA, gyrB, mreB, pyrH, recA, rpoA, topA, ftsZ). P. luminescence* was used as an outgroup. Multiple sequence alignment was performed with MEGA-X software (https://www.megasoftware.net/) (Kumar et al. 2018) using ClustalW and then MUSCLE algorithms. The primary tree was constructed with ML algorithm using using iq-tree software (Minh et al. 2020) with the best model found by iq-tree ModelFinder (GTR+F+G4); 1000 maximum likelihood bootstrap replicates were run. To estimate the time scale, the K2P substitution model (Kimura, 1980) was used. The time tree was computed using one calibration constraint according to the Pairwise Divergence times obtained with TimeTree (http://timetree.org) (Kumar et al. 2022). One taxa pair with suggested divergence time 0.016 MYA was used: *V. cholerae – V. jasicida* (Marin et al. 2017). We used complete deletion to treat missing data.

### Chemotaxonomy

For chemotaxonomic analysis, cells of type and reference strains were grown in Luria–Bertani (LB) medium (GIBCO BRL) with addition of 1.7 % NaCl under constant shaking at 120 rpm at 25°C during for 24 hours. The cells suspensions were centrifuged at 10000 g for 10 minutes.

Analysis of cellular fatty acids and polar lipids was performed as described earlier (Slobodkina et al., 2013). Respiratory quinones fraction was analyzed by reverse-phase HPLC (Jasco, Japan). Respiratory quinone standards were Q1, Q2, Q4, Q10, MK2, all from Sigma-Aldrich, along with quinone extract from *E. coli* containing Q8, MK8, DMK8. Thin-layer chromatography (TLC) standards were phosphatidylinositol, phosphatidylserine, phosphatidylcholine and phosphatidylethanolamine, all from Sigma-Aldrich.

## Results

The cells of the newly identified species in the *Vibrio* genus were isolated from coastal waters of the Black Sea (Russia) and intestines of the Black Sea horse mackerel. The cells of novel species in the *Vibrio* genus are slightly curved rods, motile, with one polar flagellum with a diameter of 1.0-1.5 µm and a length of 2.0-3.0 µm.

The strains VNB-15^T^, VNB-16 and SChm4 grow on TCBS as green colonies. On Marine agar they grow as luminescent translucent colonies. Based on the results obtained by Vitek® analyzer and other Microtests the isolated strains differ from the other types of *Vibrio*.

All three strains can produce β-glucosidase, do not form ornithine decarboxylase, acid phosphatase, N-acetyl-β-glucosaminidase and lipase. They can utilize glucose, but cannot utilize mannitol, inositol, sorbitol, arabinose, sucrose, or sodium citrate (Table 1).

**Table 1.**
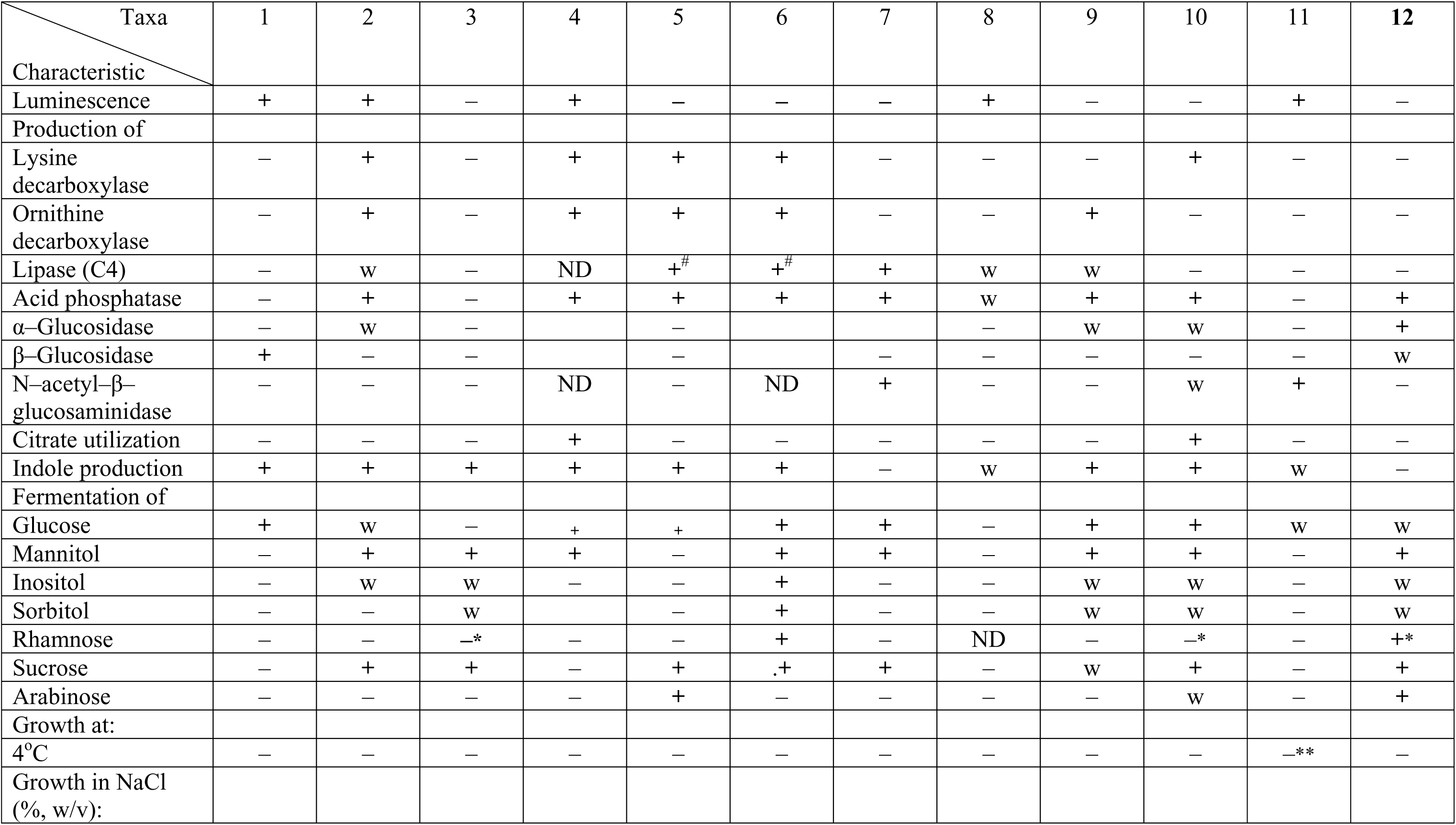
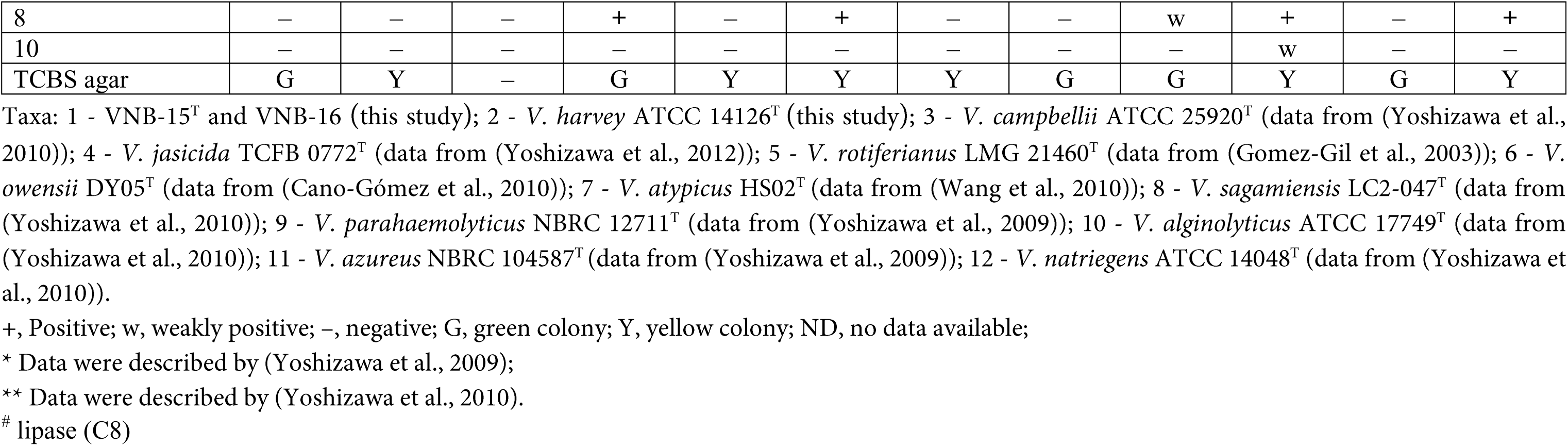
Phenotypic characteristics for differentiating the novel species *Vibrio aquamarinus* sp.nov. from phylogenetically related species of genus *Vibrio*.

Chemotaxonomic characteristics of strain VNB-15^T^ are quite similar to those of reference strain *V. harvey* ATCC 14126^T^. Major cellular fatty acids (CFA) of strain VNB-15^T^ are C16:1 (47.9 %), C16:0 (17.5 %), and C18:1ω9c (8.5 %), the same CFA are major in strain ATCC 14126^T^ (36.1 %, 12.2 % and 11.8 %, respectively) (Table S2). Respiratory quinones of strain VNB-15^T^ are composed of Q8 (67 %) and DMK-8 (33 %), whereas for the strain ATCC-14126^T^ the relative amounts of respiratory quinones are 54 % of Q8 and 46 % of DMK-8. Polar lipids of both strains are cardiolipin, phosphatidylethanolamine, phosphatidylglycerol, phosphatidylserine, unidentified aminophospholipid with Rf1 0.02-0.08 and Rf2 0.02-0.08, and unidentified polar lipid with Rf1 0.00-0.25 and Rf2 0.6-0.7, which may be detected by phosphomolybdic acid assay only (Fig S9).

Sequence data of 16S rRNA and housekeeping genes *gyrB*, *recA*, *pyrH*, *gapA, rpoA* , *mreB*, *ftsZ*, *topA* were obtained for each isolated strain (GenBank numbers in Supplementary Table S1 and Figs S1-S8). The results of phylogenetic analysis based on the comparison of the 16S rRNA genes sequences show that VNB-15^T^, VNB-16 and SChm4 strains belong to the *Vibrio* genus. These strains show phylogenetic similarity to several species of the *Vibrio* genus: about 99.56 with *V. hyugaensis*, 99.55% with *V. jasicida* (TCFB 0772^T^), 99.49% with *V. owensii* (DY05^Т^), and 99.07% with *V. sagamiensis* (LC2 - 047^T^) (Fig. 1).

**Figure 1.**
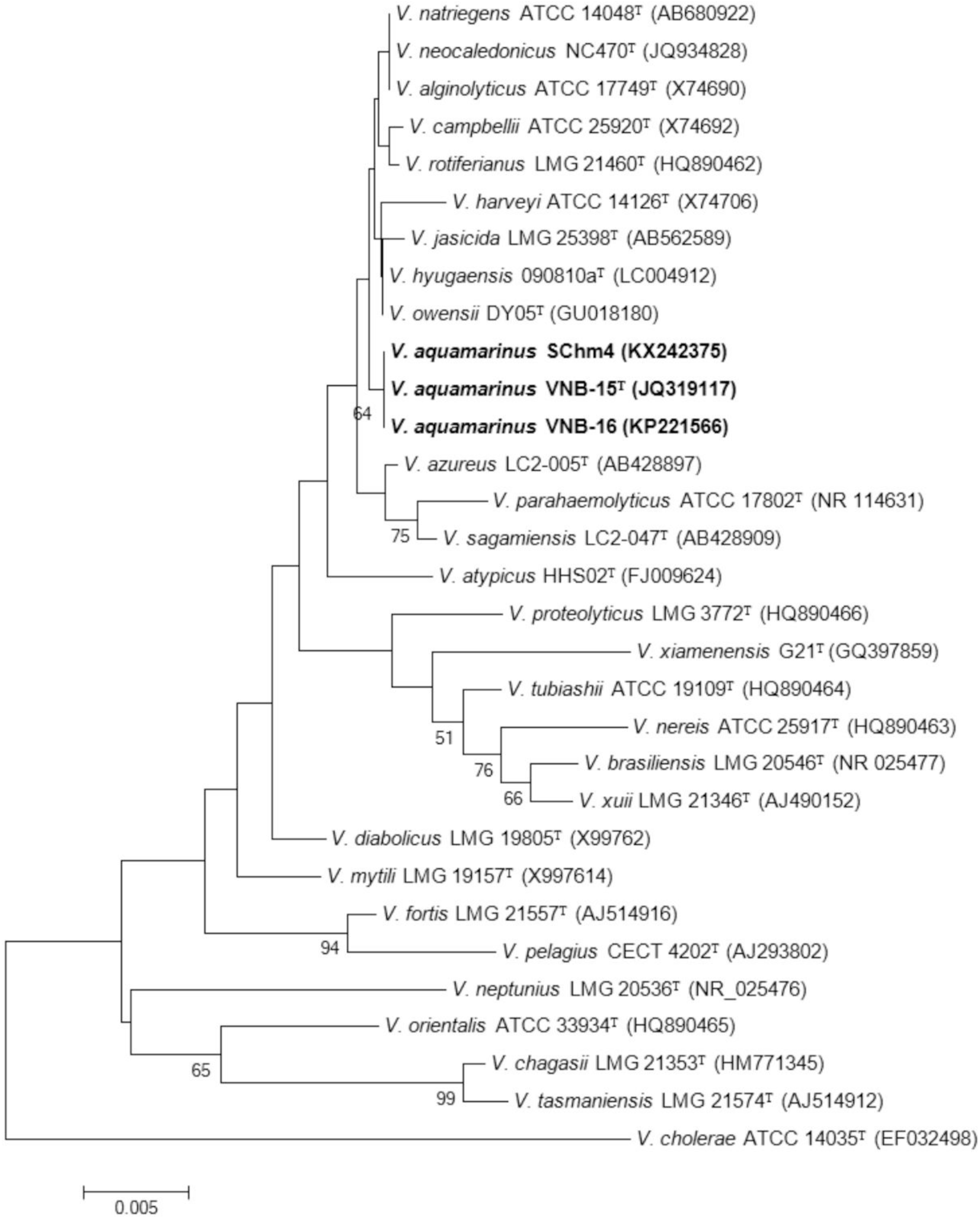
Phylogenetic tree based on the analysis of 16S rRNA sequences, constructed by the Neighbor-Joining method. Bootstrap percentages from 1000 replicates appear next to respective branches.). Out group Bar, 0.5 % estimated sequence divergence.

The combined tree constructed on the basis of seven housekeeping genes (*recA, gyrB, mreB, topA, ftsZ, pyrH, gapA*) shows that the strains VNB-15^T^, VNB-16 and Schm4 form a cluster separately from other species of the *Vibrio* genus (Fig. 2). For each of the eight housekeeping genes, phylogenetic trees are also presented in Supplementary Table S1 and Figures S1-S8. In all presented trees, the strains VNB-15^T^, VNB-16 and SChm4 co-cluster separately from other species of the *Vibrio* genus.

**Figure 2.**
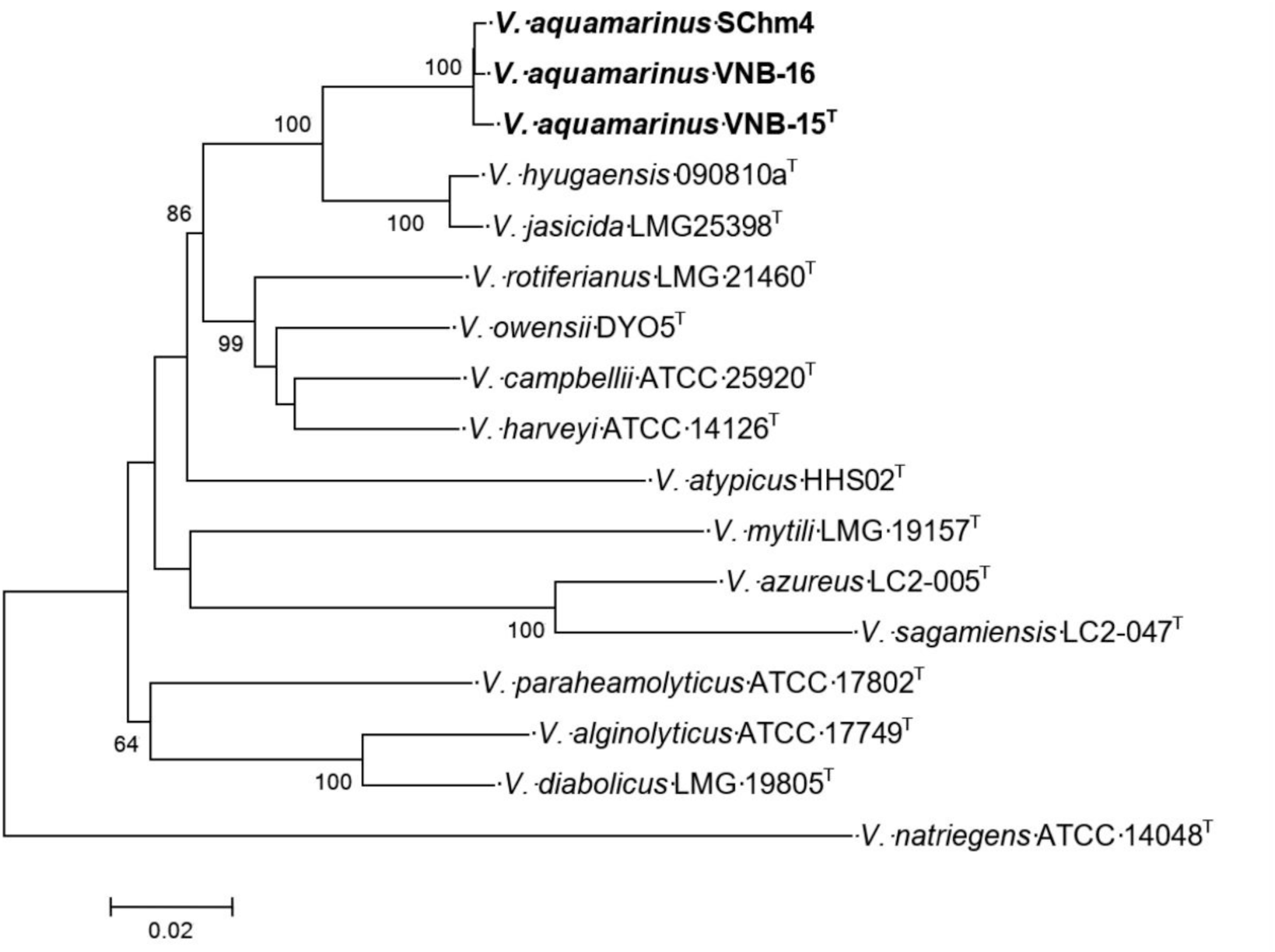
Phylogenetic tree based on the analysis of combined sequences of seven housekeeping genes *recA, gyrB, mreB, topA, ftsZ, pyrH, gapA*, for a total of 5600 bp, constructed by the Neighbor-Joining method. Bootstrap percentages from 1000 replicates appear next to the respective branches). Bar, 2% estimated sequence divergence

The genome of this bacterium was sequenced and assembled (NCBI data base SUB14585067). The total length of the assembled genome was 6230140 pairs of nucleotides, includes 2 chromosomes and a plasmid. The GC composition of the VNB-15^T^ genome was 45.14%. A phylogenetic tree was built based on this assembly and 12 complete genomes of Vibrio bacteria (Fig.3)

**Figure 3.**
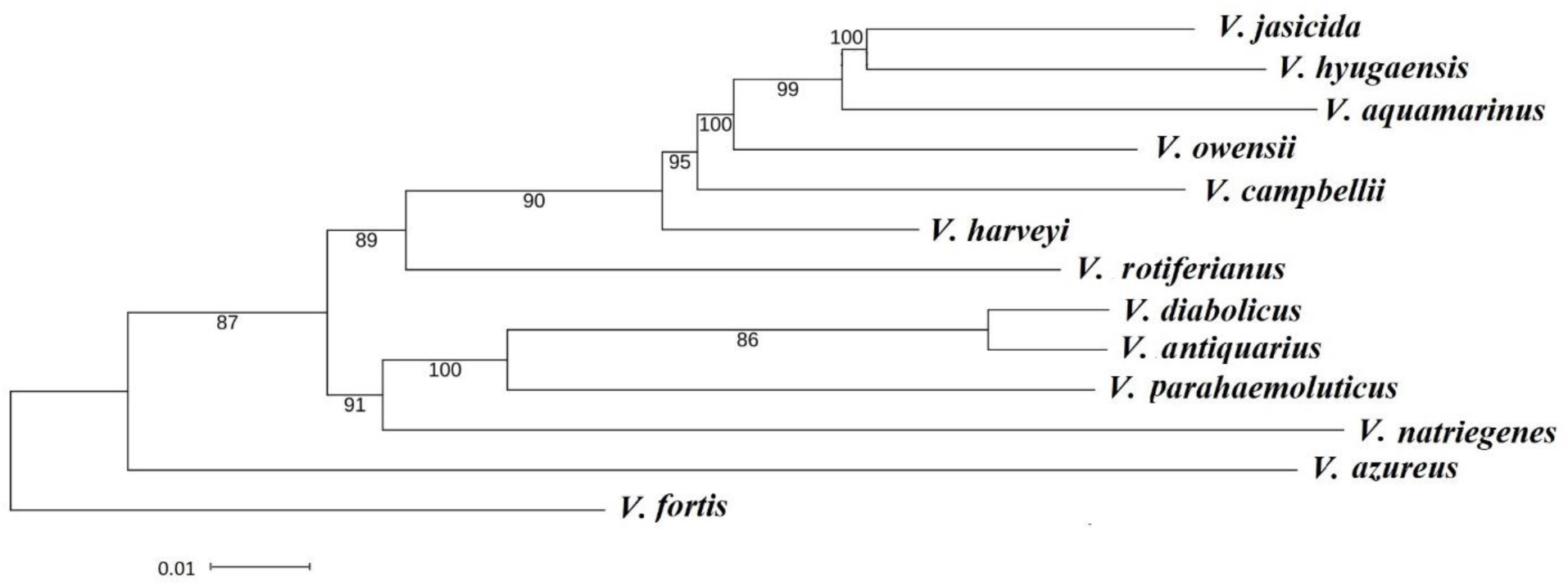
A phylogenetic Bayesian tree based on 12 complete genomes of vibrio bacteria, including *V. aquamarinus*, with a total length of 5 million base pairs. The bootstrapping percentage of 1,000 replicas is listed next to the corresponding branches.

**Figure 4.**
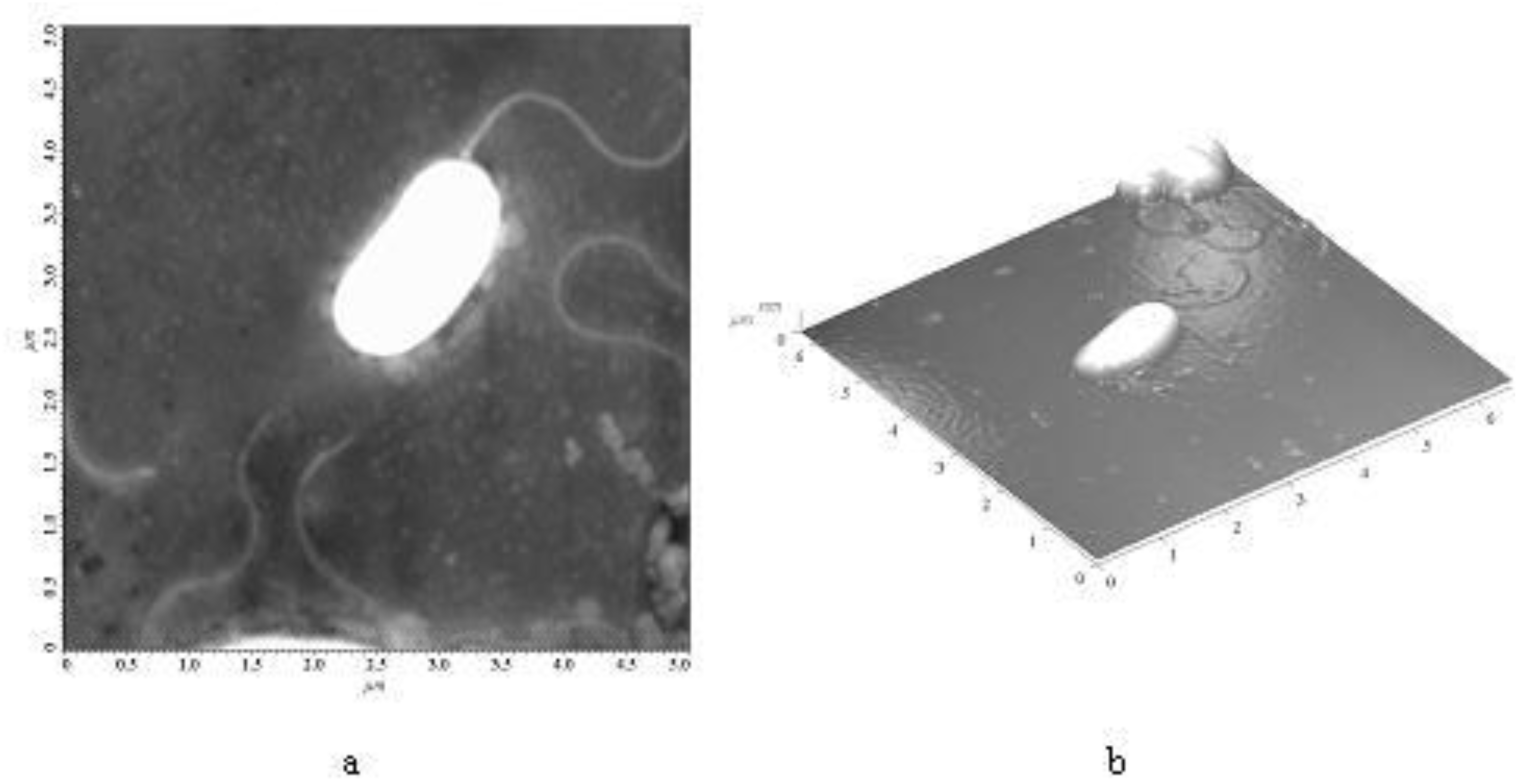
AFM 2D-image (a) and 3D-image (b) of the surface of *Vibrio aquamarinus* sp. nov.

As can be seen from the phylogenetic tree shown in Fig. 3, the sequence differences of the complete VNB-15^T^ genome from the genomes of the closest *V. hyugaensis* and *V. jasicida* are greater than those of *V. hyugaensis* and *V. jasicida* among themselves. Maximum similarity VNB-15^T^ genome is 98% with *V. hyugaensis* and 96% with *V. jasicida*.

Based on all data collected, including phylogenetic analysis, morphological, physiological and biochemical characteristics, we propose classifying strain VNB-15^T^ as a novel species in the genus, for which we suggest the name *Vibrio aquamarinus* sp. nov and the VNB-15^T^ as the type strain (VKPM B-11245^T^ = DSM 26054^T^).

Differs from closely related species by utilization of glucose, mannitol, inositol, sorbitol, rhamnose and sucrose, and by formation of lysine decarboxylase, ornithine decarboxylase, lipase, acid phosphatase, α-glucosidase, β-glucosidase, N-acetyl-β-glucosaminidase enzymes.

The luminescence spectrum of *V. aquamarinus* sp. nov. falls in the “blue” spectral range. The maximum of bioluminescence spectrum is 478 nm. *V. aquamarinus* sp. nov. are found both in the intestines of fish and in sea water.

The time tree analysis showed that the divergence of *V. aquamarinus* and *V. yasicida* strains occurred approximately 1,500 years ago (fig. S10). Because the colonization of newly formed Black Sea by fish could have taken hundreds or even thousands of years. Moreover, the comprehensive study (Bahr et al. 2006) based on High-resolution stable oxygen isotope (δ18O on ostracod shells) from Black Sea revealed that the refilling of the Black Sea and its reconnection with the Mediterranean Sea led to a gradual salinization, reaching the modern around 2000 years ago. This divergence time is entirely consistent with the hypothesis of Mediterranean fish species and their microbiota dispersal into the Black Sea.

We would also like to add that investigated strains of *V. aquamarinus* were isolated both from the sea water and from the intestines of Black Sea horse mackerel (*Trachurus mediterraneus ponticus*), genetically significantly isolated from the mackerel of the Mediterranean Sea (Slynko et al., 2018). Thus, the ecological niches of *V. aquamarinus* include both seawater and the intestines of fish, predominantly inhabiting the waters of the Black Sea). Obviously, fish intestine habitat creates a possibility for the intraspecies gene exchange in bacteria *V. aquamarinus* and, as a result, the separation of this species from the closely related forms found in the Mediterranean. It is likely that *V. aquamarinus* speciation event took place due to its relative isolation from populations of related species in the world Ocean in general and the Mediterranean in particular. This hypothesis is supported by its finding in the intestines of Black Sea horse mackerel, which harbors a variety of its strains (here we described the strain SChm4).

### Description of V. aquamarinus sp. nov

*V. aquamarinus* (a.qua.ma’ri.nus. N.L. masc. adj. *aquamarinus* aquamarine is a color that is a bluish tint of cerulean toned toward cyan, referring to the color of emitted light).

Cells are slightly curved Gram-negative rods, oxidase-positive, catalase-positive, motile with one polar flagellum, 1.0-1.5 μm wide and 2.0-3.0 μm long (Fig. 3). Colonies are rounded, with smooth margin, translucent, luminescent, have yellowish color. On TCBS agar, the colonies are green, 3-5 mm in diameter. Cells grow in either aerobic or anaerobic conditions at 10-35 °C, optimum being 20-25 °C, at 0.5-5.0 % NaCl (w / v), optimum being 1.0-4.0 %, and at pH 6.0-9.0, optimum being 7.0-8.0. Growth is supported by D-glucose, D-maltose, but not by D-cellobiose, arabinose, D-mannose, rhamnose, D-tagatose, D-trehalose, sucrose, raffinose or lactose. Adonitol, L-arabitol, D-mannitol, D-sorbitol, inositol, dulcitol, malonate, 5-keto-D-gluconate, L-lactate, succinate, L-malate, and citrate do not support growth. Strain is indole formation positive, hydrogen sulfide formation negative. It does not hydrolyze gelatin, but hydrolyzes starch. Voges-Proskauer test is negative. Strain is positive for the following enzyme activity: Ala-Phe-Pro-Arilamidase, β-glucosidase, L-proline-arilamidase, amylase, nitrate reductase. Negative for the following enzyme activity: L-pyrrolidone-arilamidase, D-cellobiose, β-galactosidase, β-N-acetyl-glucosaminidase, glyutamilarilamidase pNA, γ-glutamyl transferase, β-ksilozidase, β-alaninarilamidase pNA, lipase, palatinose, tirozinarilamidase, urease, α-glucosidase, β-N-acetyl-galaktozaminidase, α-galactosidase, phosphatase, glitsinarilamidase, ornithine decarboxylase, lysinedecarboxylase, β-glucuronidase, Glu-Gly-Arg-arilamidase, phenylalanine deaminase, arginindegidrolase, gelatinase. Not resistant to O/129 (2,4-diamino-6,7-diisopropylpteridine) vibriostatic agent. Major cellular fatty acids are C16:1 (47.9 %), C16:0 (17.5 %), and C18:1ω9c (8.5 %), respiratory quinones are Q8 (67 %) and DMK8 (33 %).

The type strain is VNB-15^T^ (=VKPM B-11245^T^ = DSM 26054^T^). Reference strains are VNB-16 (=VKPM В-12136), SChm4 (=VKPM B-14175)

The type strain VNB-15^T^ was isolated from the Black Sea near the town of Abrau-Durso, Russia.

## Acknowledgments

Isolation of VNB-15 and VNB-16 strains and phenotypic analysis were supported by the Ministry of Science and Higher Education of the Russian Federation within the framework of the state assignment in the field of scientific activity (project no. FENW-2024–0026). Isolation of SChm4 strain, assembly and annotation of the complete genome VNB15^T^, analysis of the phylogeny were funded by RFBR project № 22-14-00124-П. The work of A.A.N. at the Gubkin University (lipid extraction and analysis) was supported by the Russian Science Foundation, project 20-79-10388.

The authors are grateful to Pechenyi A.P. for the help with copyediting.

## Author contribution

M.A.S - isolation of the type strains VNB15^T^ and VNB16

S.A.Khr. - DNA extraction, sequencing and analysis of the phylogeny

I.S.S. – Phenotypic analysis of isolated strains

E.V.M. – assembly and annotation of the complete genome VNB15^T^, analysis of the phylogeny

A.A.N.– analysis of cellular fatty acids and polar lipids

K.M.N. – preparing the manuscript

S.V.B. – preparing the manuscript

S.M.R. – sequencing

K.A.A. – sequencing

V.B.Sh.– production of micrographs (the morphology of cells and their flagella)

V.A. Ch – sequencing

I.V. Manukhov - isolation of SChm4 strain from fish guts, analysis of the phylogeny, summarily data analysis and preparing the manuscript

All authors edited and approved the manuscript

